# Codon Deoptimization of Multispecific Biologics Reduces Mispairing During Transient Mammalian Protein Expression

**DOI:** 10.64898/2026.01.05.694717

**Authors:** Timothy Z. Chang, Weijun Ma, Jane Guo, Jiali Hu, Kalie Mix, Yi Tang, Karen Wong, Eva Bric-Furlong, Amanda Lennon, Brian Hall, Dietmar Hoffmann

## Abstract

Codon optimization is utilized in biologics design to maximize protein expression. Selecting the host organism’s most frequently used codons for each amino acid can significantly enhance recombinant protein expression yields. However, non-optimal codons in mRNA can be critical for functional protein production through inducing pauses in or attenuating protein translation. In our study, we have investigated the effect of deoptimizing serine codons in biologics by shifting them from the five most frequently used codons to the least (TCG).

Rare serine codons were strategically inserted into the coding sequences of the constant regions in a trispecific antibody (Protein 1), a bispecific antibody (Protein 2), and multiple non-proprietary bispecific antibodies. We observed that inserting 1-2 rare serine codons within an open reading frame led to expression changes that reduced the formation of mispaired 2x light chain and half-molecule species. Protein purity was drastically increased by incorporating two deoptimized serine codons into a single chain. Notably, we observed a negative correlation between total protein expression yield and final product purity.

Taken together, our work demonstrates that incorporation of deoptimized serine codons into a single chain can significantly influence multispecific biologic pairing and enhance final product purity. Our findings align with existing literature showing that rare codon usage modulates translation kinetics and protein folding. Future investigation is warranted to enable *a priori* identification of the rate-limiting chain in multispecific biologics, thereby guiding strategic codon deoptimization prior to expression.

## 1 Introduction

Bispecific and multispecific therapeutics are emerging as increasingly important biotherapeutic modalities. The number of clinical trials continues to grow year after year, with over 300 trials initiated in 2024 alone (1). This rising therapeutic demand for multiple target engagement has led to a need for robust recombinant protein design and expression strategies. To meet these challenges, numerous novel molecular configurations have been engineered to simultaneously engage multiple targets and couple them with appropriate Fc-mediated immunoeffector functions (2).

A significant challenge in multispecific biologic production is mispaired impurity formations. As most biologics are composed of one or more pairs of heavy and light chains, incorrect pairing can hinder successful recombinant expression of multi-chain molecules. Various strategies have been developed to enhance correct molecule pairing, including knob-and-hole mutations to promote properly paired half-molecules (3), and Fab interface mutations to facilitate correct heavy/light chain pairing (4, 5). While these mutations can improve chain association, they may also adversely affect protein expression and stability. An alternative approach is the use of a common light chain on both halves of a bispecific molecule, which can sidestep the need for such engineering altogether (6).

However, even with chain-chain interface mutations designed to drive proper chain association, mispairing can still occur. One potential reason for this is mismatched chain expression levels, which can cause stoichiometry-driven mispairing. Two common approaches to balancing chain expression levels involves adjusting the ratio of transfected plasmids encoding each of the chains (7) or using different promoters or other genetic elements to fine-tune mRNA transcription (8).

An understudied factor in protein expression is codon usage, which can more precisely balance chain expression. Codon optimization is an essential part of recombinant biologics design. It is commonly performed during the reformatting of sequences identified in the discovery phase into therapeutic candidate molecules designed for overexpression in mammalian cells (9). Generally, this process involves converting the codons of the discovered sequence into those used at a frequency similar to that natively found in the host organism - *Homo sapiens* for Expi293 and *Cricetulus griseus* for CHO cell lines. By using the most frequently used codons in each organism for recombinant protein expression, codon optimization aims to facilitate the rapid translation of mRNA, preventing translational stalling and minimizing nascent polypeptide misfolding or degradation (10). In a single molecule study, codon optimization was observed to increase the mRNA translation rate in mammalian cells from approximately 3.1 to 4.9 codons/second (11).

Codon <u>de</u>optimization, the deliberate incorporation of rare codons into a recombinant coding sequence, may be an alternate design strategy. Early evidence linking secondary structure boundaries to differences in codon usage led researchers to suspect that codon frequency plays a role in co-translational folding(12, 13). Additionally, silent mutations in the E*. coli* chloramphenicol acetyltransferase from rare codons to common ones were found to increase both protein yield and misfolding (14), highlighting the role of rare codons in regulating functional protein production.

These findings have found practical application through the strategic alteration of codon usage in viruses to generate live-attenuated vaccines (15, 16). Reduced translation efficiency caused by non-optimal codon usage reduces viral replication efficiency and pathogenicity, while still enabling a robust anti-viral cellular immune response(17). Although non-optimal codon usage can stall translation and lead to reduced expression via cotranslational ubiquitination and other pathways(18, 19), it can also improve expression of difficult to fold proteins (20, 21). In particular, rare codons can induce translational pausing in the nascent polypetide chain, allowing more time for co-translational folding and promoting the formation of properly conformed proteins (22).

This principle has also been explored for enhancing yield of biologics. Kumada et al. employed rare serine codons in a single chain variable fragment (scFv) (G_4_S)_3_ linker to enhance expression in *E. coli* (23). In mammalian cell-based expression, Magistrelli et al. introduced non-optimal codons into a lambda light chain to improve expression of a bispecific κλbody by enhancing kappa light chain expression (24). These studies raise two possible mechanisms by which codon deoptimization can enhance biologics production. First, incorporating non-optimal codons into an overexpressed chain may attenuate its expression, promoting balanced chain production. Second, incorporating non-optimal codons into an underexpressed chain may stabilize it through improved co-translational folding, thereby increasing its production and balancing chain expression.

In this work, we investigated the impact of codon deoptimization on the expression of multispecific biologics that exhibit significant mispairing issues. Using high throughput transient mammalian protein expression and characterization, we systematically evaluated various combinations of optimized and deoptimized chains across multiple bi- and tri-specific protein formats. We explored the influence of codon deoptimization position and assessed whether modifying one or multiple chains could improve both the yield and structural integrity of the expressed products. Our findings provide new insights into codon usage as a tunable parameter which may enhance the quality and balance of complex biologic therapeutics.

## 2 Materials and Methods

### 2.1 DNA Design and Preparation

Nucleic acid sequences were designed in Snapgene (Dotmatics, Boston, MA) and ordered as double-stranded gene fragments (Integrated DNA Technologies, Coralville, IA). Sequences were cloned into the pTT5 expression vector (National Research Council, Ottawa, ON) using the SnapInfusion DNA assembly kit (Takara Bio USA, San Jose, CA). Assembled plasmids were transformed into chemically competent Mix&Go® *E. coli* DH5α according to the manufacturer’s instructions (Zymo Research, Irvine, CA). Overnight cultures were grown in Plasmid Plus® broth (Thomson, Oceanside, CA) with 100 µg/mL carbenicillin. Plasmids were purified by the Qiagen Miniprep kit (Qiagen, Germantown, MD), quantified by absorbance at 280 nm and/or by Qubit® fluorometric quantitation (Thermo Fisher Scientific, Waltham, MA). Plasmids were sequenced with Sanger and/or whole-plasmid sequencing (Azenta, Burlington, MA) and verified in Snapgene and Genedata Biologics (Genedata, Lexington, MA).

### 2.2 Cell Culture

Engineered plasmids were transiently transfected into Expi293 cells and cultured according to manufacturer’s instructions (Thermo Fisher Scientific, Waltham, MA). Briefly, 2mL Expi293 at 3*10^6^ cells/mL were aliquoted into 24 well round-bottom plates (Thomson, Oceanside, CA) and transfected with 2µg plasmid DNA according to the Expifectamine transfection protocol (Thermo Fisher Scientific, Waltham, MA). Cells were grown at 37°C with 8% CO_2_ and 80% relative humidity shaking at 240 rpm in a 25mm orbital shaker. Recombinant proteins were harvested from cell culture supernatants at 5 days post-transfection.

### 2.3 High-Throughput Protein Expression

The fully automated mammalian cell secretory overexpression system, Protein Expression and Purification Platform (PEPP; GNF, San Diego, CA, USA), was used to express and to purify the bispecific antibody molecules. In brief, 35 mL of cultures seeded with 3 × 10^6^ Expi293F cells (Thermo Fisher Scientific A14527, Waltham, MA, USA) were transfected with 34 μg of plasmid DNA using the Expifectamine 293 transfection reagent per the manufacturer’s protocol (Life Technologies Corporation, Carlsbad, CA, USA). Cells were incubated at 37°C in an 8% CO_2_ environment for 5 days. Culture supernatants were harvested and applied to MabSelect SuRe protein A resin (GE Healthcare 17-5438-02, Chicago, IL, USA) for gravity flow purification. Proteins were desalted using Nap-10 Sephadex columns (GE Healthcare 17-0854-02) and eluted with 1X Dulbecco’s Phosphate-Buffered Saline (Thermo Fisher Scientific 14190136), resulting in a final storage solution at pH 7.4 for each protein sample.

### 2.4 Protein Analysis

Cell culture supernatants were collected by brief centrifugation and analyzed by SDS-PAGE and Western Blotting. Briefly, 60 µL sample was mixed with either 20 µL 4x NuPAGE™ LDS (Thermo Fisher Scientific, Waltham, MA) or 60 µL Laemmli (Bio-Rad, Hercules, CA) sample loading dye with or without 50 mM dithiothreitol as a reducing agent. Samples were heated to 98°C for 5 minutes and run on NuPAGE™ 4-12% Bis-Tris or 4-20% Criterion^TM^ TGX Stain-Free^TM^ Precast gel (Bio-Rad, Hercules, CA) according to the manufacturer’s protocol. Stain-free gels were run using a Criterion^TM^ Electrophoresis Cell (Bio-Rad, Hercules, CA) with a Bio-Rad PowerPac Basic (Bio-Rad, Hercules, CA) at 300V, 400mA constant, for 25 minutes. Results were analyzed using ImageLab software (Bio-Rad, Hercules, CA). Stain-free bands were annotated using the LC-MS results to identify the protein of interest and the mispaired molecules present in each sample.

Capillary gel electrophoresis was used to assess protein A-purified purified protein concentration, molecular weight, and purity in high throughput. Protein Characterization was performed with the LabChip® GX Touch system (Revvity, Waltham, MA) using the Protein Express Chip (#760528) and Protein Express Assay Reagents kit (# CLS960008). Samples were processed according to the manufacturer’s protocol with the following modifications: 2 µL of 0.04 – 2 mg/mL protein sample was added to either 18 µL of Protein Express buffer (non-reducing conditions) or 18 µL of a 700 µL Protein Express Sample Buffer with 24.5 µL β-mercaptoethanol master mix(reducing conditions). 15 µL of 10x diluted ladder was used as a standard.

### 2.5 Analytical Cation Exchange Chromatography Analysis

Analytical Cation Exchange Chromatography (aCEX) analysis was performed using an Agilent 1290 Infinity II LC System (Agilent Technologies, Santa Clara, CA) on a ProPac WCX-10, 10 µm, 4 × 100 mm column (Fisher Scientific, Atlanta, GA). Protein was eluted over a 20-minute salt gradient from 0-1M NaCl (40mM MES, pH 5.6). Results were analyzed using OpenLab ChemStation software (Agilent Technologies, Santa Clara, CA).

### 2.6 Liquid Chromatography and Mass Spectrometry

2 µg of each sample was injected onto an Agilent 1290 HPLC instrument equipped with a PLRP-S column (Agilent: PL-1912-1502) operating at a flow rate of 0.3 mL/min. Protein was separated using a linear gradient of water and acetonitrile, each with 0.1% formic acid, and injected into the Agilent 6545XT Q-TOF. Source conditions are as follows: gas temp 350 °C, drying gas 12 L/min, nebulizer pressure 35 psi, sheath gas temp 400 °C, sheath gas flow 11 L/min, Vcap 400 V, nozzle volage 2000 V, fragmentor voltage 180 V, and skimmer voltage 65 V. Data was deconvoluted and analyzed using Protein Metrics Byos software.

## 3 Results

### 3.1 Transient Expression of Multispecifics Generates Mispaired Impurities

Proper pairing of heavy and light chains is critical in multispecific biologics containing three or more chains. Despite the use of engineered mutations to facilitate correct pairing (e.g. knob-in-hole, charge-pair mutations), mispaired species are still produced to a certain extent. We demonstrated this with two proteins – Protein 1 is a tri-specific, crossover dual variable region (CODV) antibody containing two tandem variable regions on one side and one variable region on the other (Figure 1A)(25). The four chains required to assemble this molecule differ in size and are resolvable by reduced SDS-PAGE. A 2x monoclonal light chain impurity is clearly visible by non-reduced SDS-PAGE with size smaller than the properly paired molecule (Figure 1B). Quantification by SDS-PAGE band intensity revealed that the protein consists of 72% intact molecule and 27% mispaired impurity (Figure 1C).

**Figure 1.**
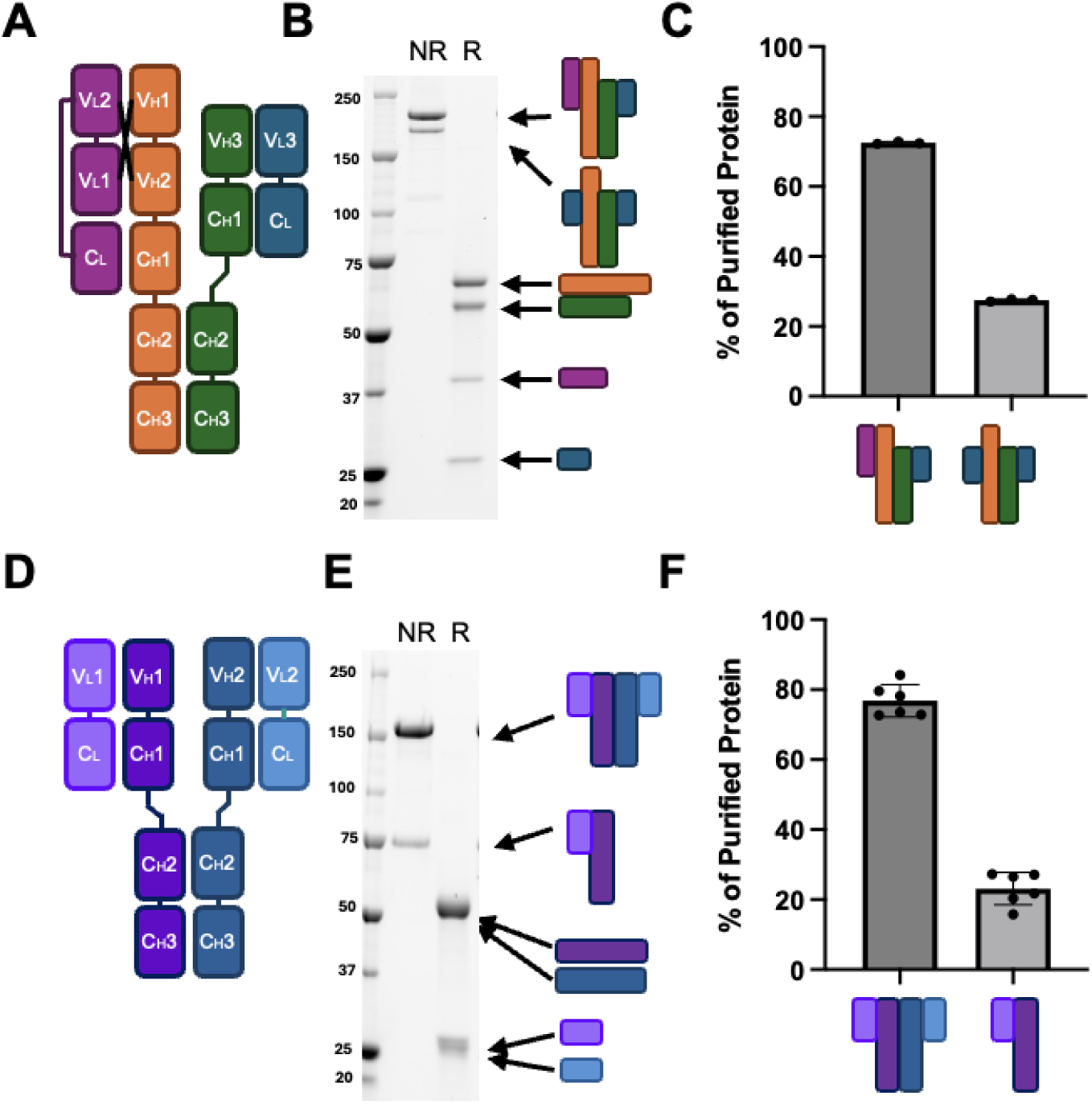
Representative mispairing issues among multispecific proteins. (A) Diagram of representative CODV antibody Protein 1 with (B) SDS-PAGE of purified material showing non-reduced component chains (NR, first lane) and reduced purified species (R, second lane) (C) Quantified SDS-PAGE bands of various impurities in Protein 1. (D) Diagram of representative bispecific antibody Protein 2 (E) Representative SDS-PAGE of purified material showing non-reduced component chains (first lane) and reduced purified species (second lane) (F) Quantified SDS-PAGE bands of various impurities in bispecific Protein 2.

A similar analysis was performed for Protein 2, a bispecific antibody consisting of a single variable region on each half of the molecule (Figure 1D). Since both heavy and light chains are roughly the same size, they cannot be resolved by reduced SDS-PAGE. However, a significant half-molecule impurity is detectable by non-reduced SDS-PAGE (Figure 1E), comprising approximately 77% of the purified material (Figure 1F). Both SDS-PAGE-based protein impurity quantifications were confirmed by mass spectrometry (Figure S1).

### 3.2 Deoptimization of Serine Codons in One Chain Reduces CODV Trispecific Mispairing

Codon deoptimization was performed by mutating one or more serine codons from the five commonly used ones to the rarest, TCG (Figure 2A). We hypothesized that altering the translation rate of one of the four chains in Protein 1 through codon deoptimization would reduce mispairing. To enable broader applicability to other biologics, we chose serine residues in the signal peptide and constant regions of all four chains for deoptimization in a screening approach (Figure 2A, black boxes, Figure 2C).

**Figure 2.**
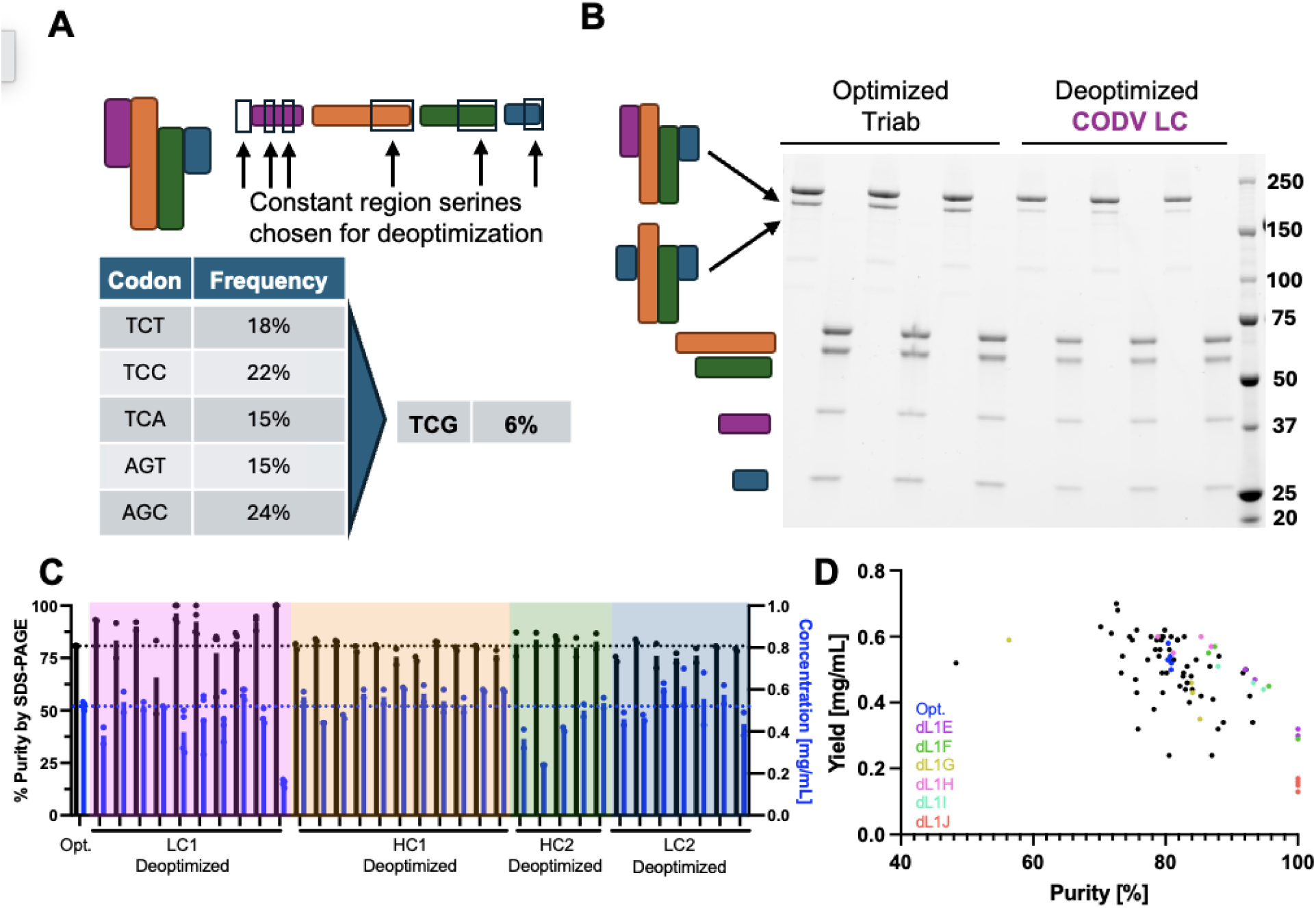
Screening constant region serines for codon deoptimzation. (A) Diagram of Protein 1 (top left) and location of mutations in various chains (top right). Frequency of common (left) and rare (right) serine codons in the human genome. (B) SDS-PAGE of three replicate non-reduced and reduced expressions of Protein 1 without (left) and with (right) deoptimized serine codons. (C) Protein purity as determined by capillary gel electrophoresis band quantitation (black bars) and purified protein yield determined by A_280_ (blue bars) for each of the deoptimized expressions performed. Colored box indicates deoptimized codon was introduced into the corresponding colored chain in panel (A). (D) Plot of individual sample purities and yields, with representative samples from deoptìmized LC1 indicated by color.

We found that deoptimizing certain serine codons in the CODV light chain (LC) (pink, Figure 2A) led to increased final product purity (Figure 2B). The increase in purity was only observed when deoptimized serine codons were incorporated into the CODV LC (Figure 2C, black bars, pink rectangle) while deoptimizing serine codons in other chains did not significantly increase purity. Notably, in the deoptimized constructs that enhanced final product purity, we observed a corresponding decrease in final purified product titer (Figure 2C, blue bars). Plotting yield against purity for each expression revealed a negative correlation between the two (Figure 2D), with deoptimized serine codons in the intra-variable region linker yielding the highest purity for a given protein yield (colored dots, Figure 2D).

These proteins made with deoptimized CODV light chains, designated dL1E through dL1J, were further characterized (Figure 3A). LC-MS analysis confirmed that deoptimizing the CODV light chain increased intact molecule purity and reduced the mAb light chain (2xLC2) impurity (Figure 3B). SDS-PAGE analysis of conditioned media (CM) harvested prior to purification showed a strong increase in CODV light chain production from the deoptimized CODV light chain sequences compared to a fully codon-optimized version (pink bar, Figure 3C).

**Figure 3.**
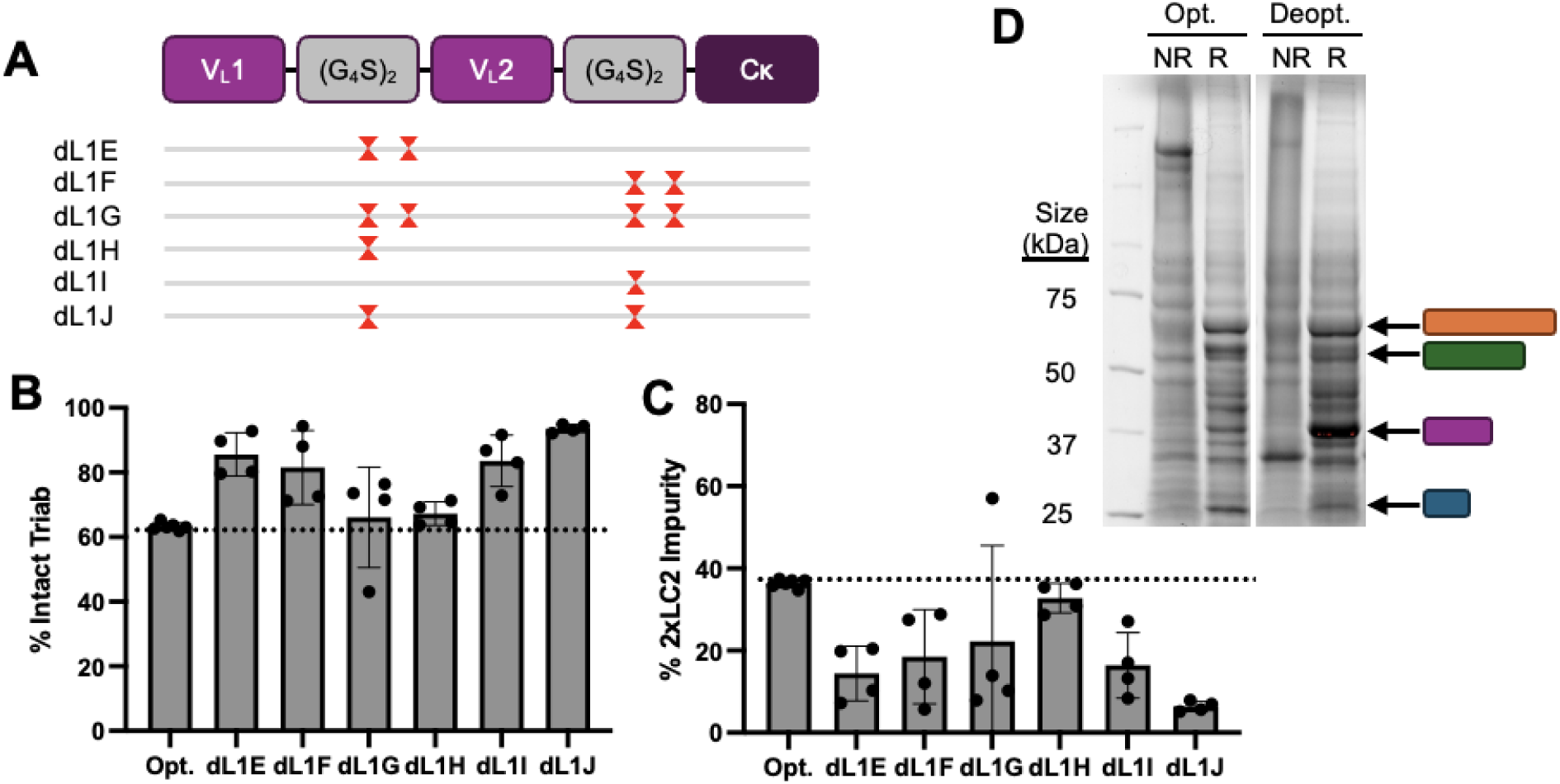
(A) Selected deoptimized LC1 plasmids showing positions of deoptimized serine(s) (red hourglass) in relation to the coding sequence of the chain (top). (B) Purity analysis by mass spectrometry of intact triab (left) and (C) 2xLC2 impurity species (right) in native and deoptimized triabs. (D) Reduced SDS-PAGE of cell culture supernatant from codon optimized (left) and a representative codon deoptimized (dL1J) (right) expressions.

### 3.4 Bispecific Protein Deoptimization Reduces Half-Molecule Impurity

A similar serine deoptimization screen was performed on Protein 2, a Y-shaped bispecific protein (Figure 4A). Given that productive serine deoptimizations in protein 1 appeared to depend more on the individual chain rather than on the specific position within that chain, the positions of deoptimized serine codons were restricted to the first two constant region serines of each chain, testing one or two deoptimizations per condition. The optimal combination for reducing mispaired impurities was found to be two deoptimized serine codons, with one in light chain 2 and one in heavy chain 2 (Figure 4B). This combination drastically reduced the amount of half-molecule impurity as shown by SDS-PAGE (Figure 4C) LC-MS (Figure 4D), and aCEX (Figure 4E). However, differences in chain expression in the CM were not distinguishable by SDS-PAGE (Figure S2).

**Figure 4.**
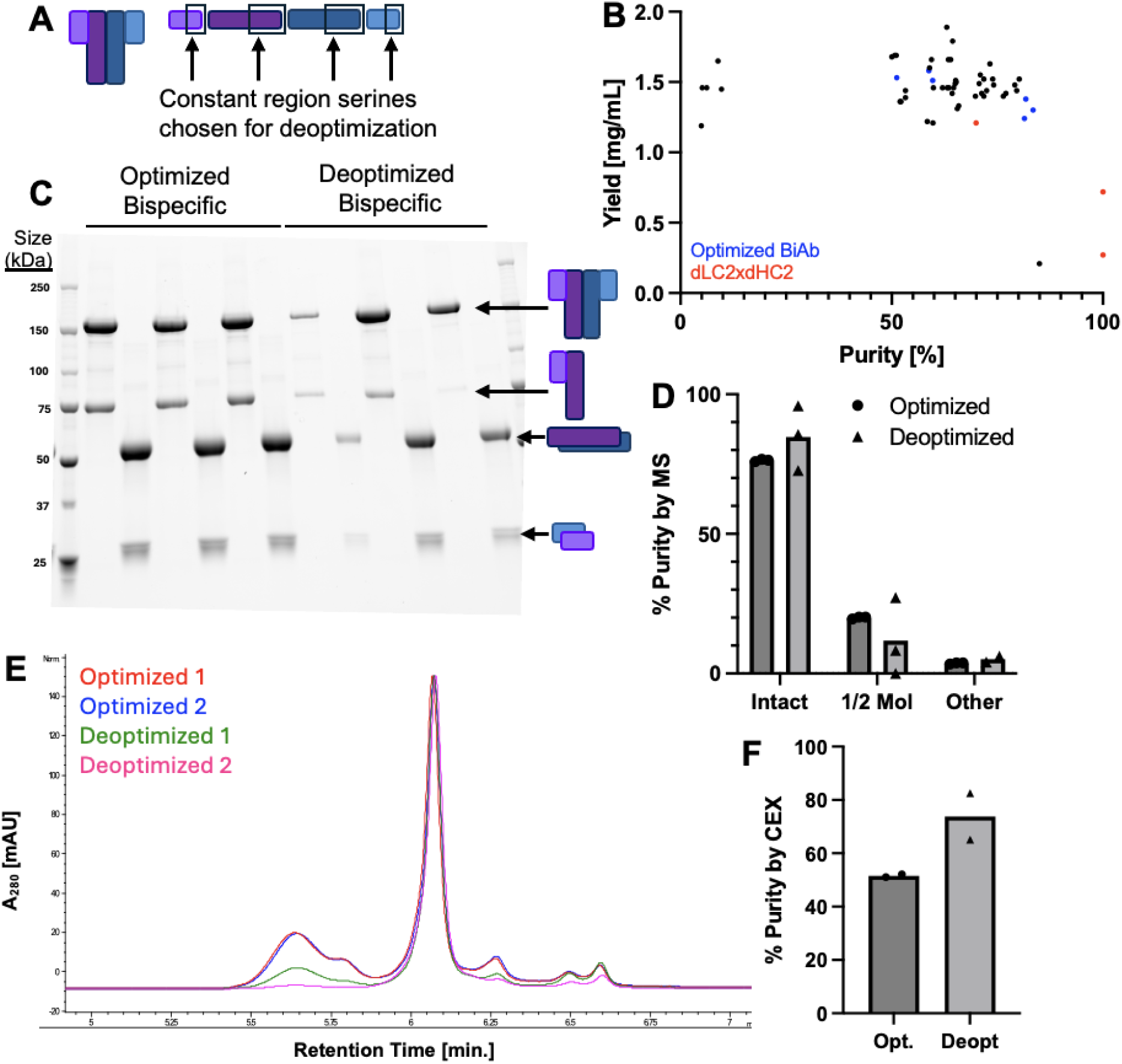
Introducing constant region deoptimized serines into a bispecific antibody. (A) Diagram of representative bispecific antibody Protein 2 (left) and location of mutations in various chains (right). (B) Plot of individual sample purities and yields, with representative samples indicated by color. (C) Three replicates of non-reduced (left column) and reduced (right column) purified samples of optimized and deoptimized Protein 2. (D) Composition of optimized and deoptimized proteins as measured by mass spectrometry. (E) Representative cation exchange (CEX) chromatographs of optimized (red, blue) and deoptimized (green, pink) proteins. (F) Quantified peak of interest purity by CEX.

Building on this strategy, we applied serine deoptimization to other generic Y-shaped bispecific proteins. One such molecule, an Adalimumab x Alemtuzumab (Figure 5A) bispecific molecule has an overexpressed Adalimumab heavy and light chain, resulting in 2x Adalimumab light chain impurities and Adalimumab half-molecule impurities (Figure 5B and 5D). Introducing two deoptimized serine codons at the start of the underexpressed Alemtuzumab heavy and light chain constant regions (Figure 5C) led to a reduction in the 2x Adalimumab LC impurity, but did not affect the Adalimumab half-molecule impurity (Figure 5D,F).

**Figure 5.**
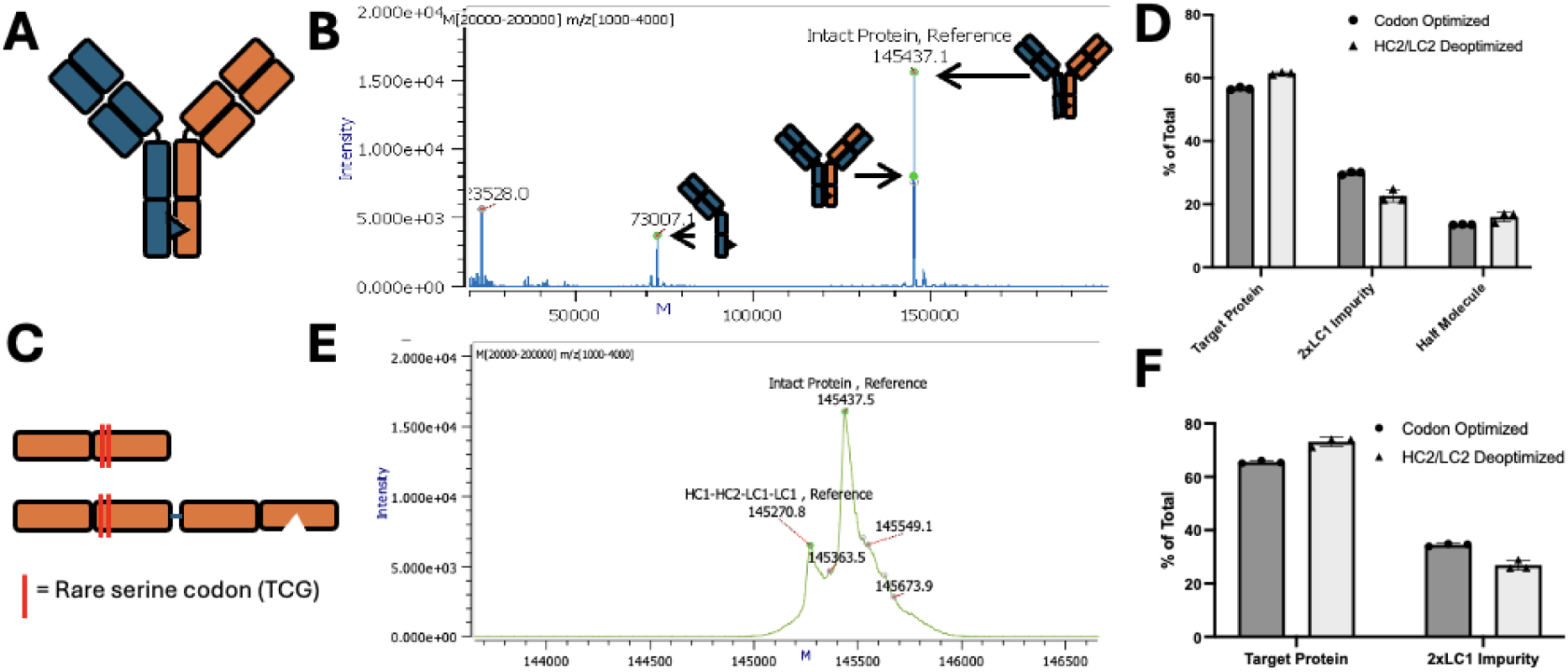
Deoptimizing an Adalimumab x Alemtuzumab bispecific reduces 2xLC impurity. **(A)** Diagram of adalimumab x alemtuzumab knob-in-hole bispecific. **(B)** Mass spectrum of codon-optimized adalimumab x alemtuzumab expression. Cartoons of various species produced shown below. **(C)** Diagram of alemtuzumab side of the bispecific molecule showing locations of deoptimized serine codons introduced in light (top) and heavy (bottom) chains. **(D)** Percentage of total purified protein from codon optimized (dark grey) and codon deoptimized (light grey) molecules. **(E)** Alternative analysis of mass spectra (top) comparing only paired species. **(F)** Percentage composition of each species in alternative analysis.

## 4 Discussion

Mispaired impurities are a common challenge in the production of bispecific and multispecific biologics,. The additional purification steps needed to generate final, high purity product can increase timelines and resource usage during biotherapeutic research and downstream process development (26). In this study, we explored whether codon deoptimization, the deliberate introduction of rare codons into a protein coding sequence, could be used to reduce impurities by modulating the expression kinetics of individual chains in several types of multispecific biologics.

Our results demonstrate that targeted serine codon deoptimization, specifically the substitution of commonly used serine codons with the rare TCG codon, can significantly improve the purity of certain multispecific proteins during transient expression. In the case of a representative CODV trispecific, deoptimization of the CODV light chain with two serine codons increased final purity from approximately 70% to around 95%. A similar purity increase from 75% to approximately 85% was observed for bispecific Protein 2. These improvements were associated with increased expression of the underrepresented chain, suggesting that codon deoptimization can be used not only to attenuate overexpressed chains (24) —as previously reported—but also to boost expression of underexpressed chains, thereby improving chain pairing and final product quality.

A key observation was the inverse relationship between product purity and yield. While deoptimization improved purity, it often reduced overall protein yield. (Figure 2D, 4B). This tradeoff may still be advantageous, as a higher initial purity can reduce the need for additional, yield reducing purification steps (26). Our findings are consistent with those of Komar et al., who showed a similar tradeoff between expressed protein quantity and quality resulting from different codon optimizations(14). Taken together, our observation of increased production of the deoptimized chain in the CM (Figure 3C) suggests that protein production is limited by the synthesis of the slowest-folding or least-expressed chain, and that adjusting the translation rate of this rate-limiting chain can shift the balance of productive assembly versus mispairing– i.e. increasing translation and/or folding of the rate-limiting light chain 1 while decreasing translation of the other chains required for the molecule, resulting in less total protein produced.

It is also possible that certain mispairings cannot be ameliorated by codon deoptimization, as evidenced by the inability of deoptimized Alemtuzumab chains to reduce the Adalimumab half-molecule impurity in the Adalimumab x Alemtuzumab bispecific (Figure 5D).

Taken together, our findings highlight the need for strategic placement of rare codons, ideally informed by an understanding of which chain is rate-limiting in a given multispecific format. Current codon optimization algorithms typically avoid rare codons altogether, potentially missing opportunities to fine-tune expression and folding. Notably, in synthetic fusion proteins like CODV LCs, inserting rare codons into flexible linkers or structurally complex regions may help synchronize folding and assembly for many. There is a correlation between rare codons, reduced translation speed and co-translational folding. However, predicting folding bottlenecks from sequence alone remains a significant challenge. In the absence of computational algorithms for identifying preferred codon usage for each chain in a given multispecific, pilot expression data on individual chains may provide insight into expression challenges associated with different components of a multispecific biologic. In the case of synthetic fusion proteins like the CODV LC of Protein 1, it is also intuitive to incorporate rare serines into GS linkers between domains that are not naturally found together, as these are more likely to suffer from folding difficulties.

## 5 Conclusion

Overall, we have demonstrated that incorporating deoptimized serine codons into the constant regions of some antibody-based biologics can improve their expression purity during transient mammalian expression. However, we also identified a tradeoff between purity and yield, highlighting the need for careful optimization, and the success of this strategy depends on the specific chain interactions and folding dynamics of each multispecific molecule. When designing biologics, codon optimization and deoptimization should be used strategically to maximize the final product’s yield and purity. When applied judiciously, they offer a powerful means to balance productivity and purity, accelerating the development of high-quality multispecific therapeutics.

## 7 Supplemental Figures

**Figure S1.**
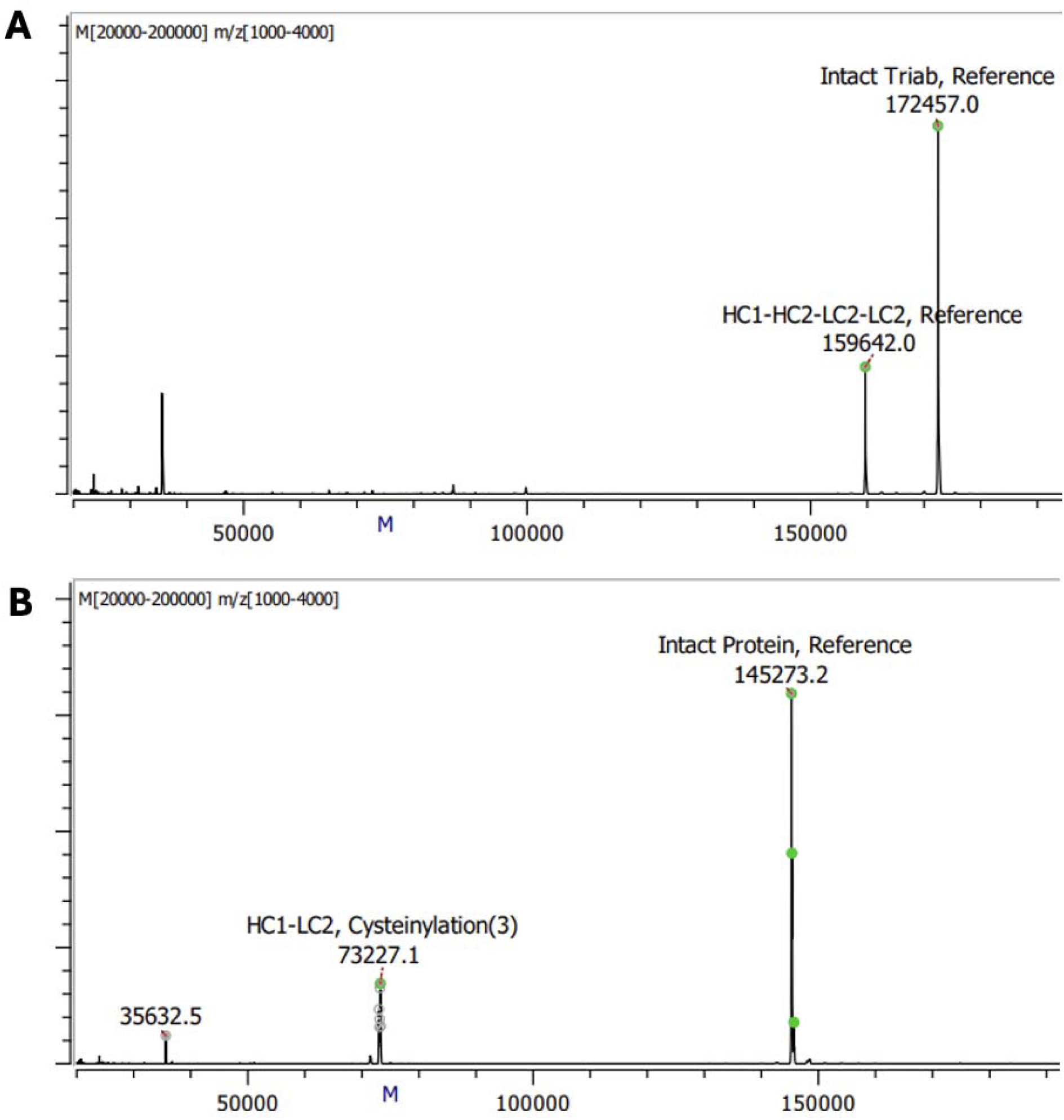
Representative mass spectra of the CODV trispecific Protein 1 (A) and the bispecific protein (B) with their associated impurity peaks.

**Figure S2.**
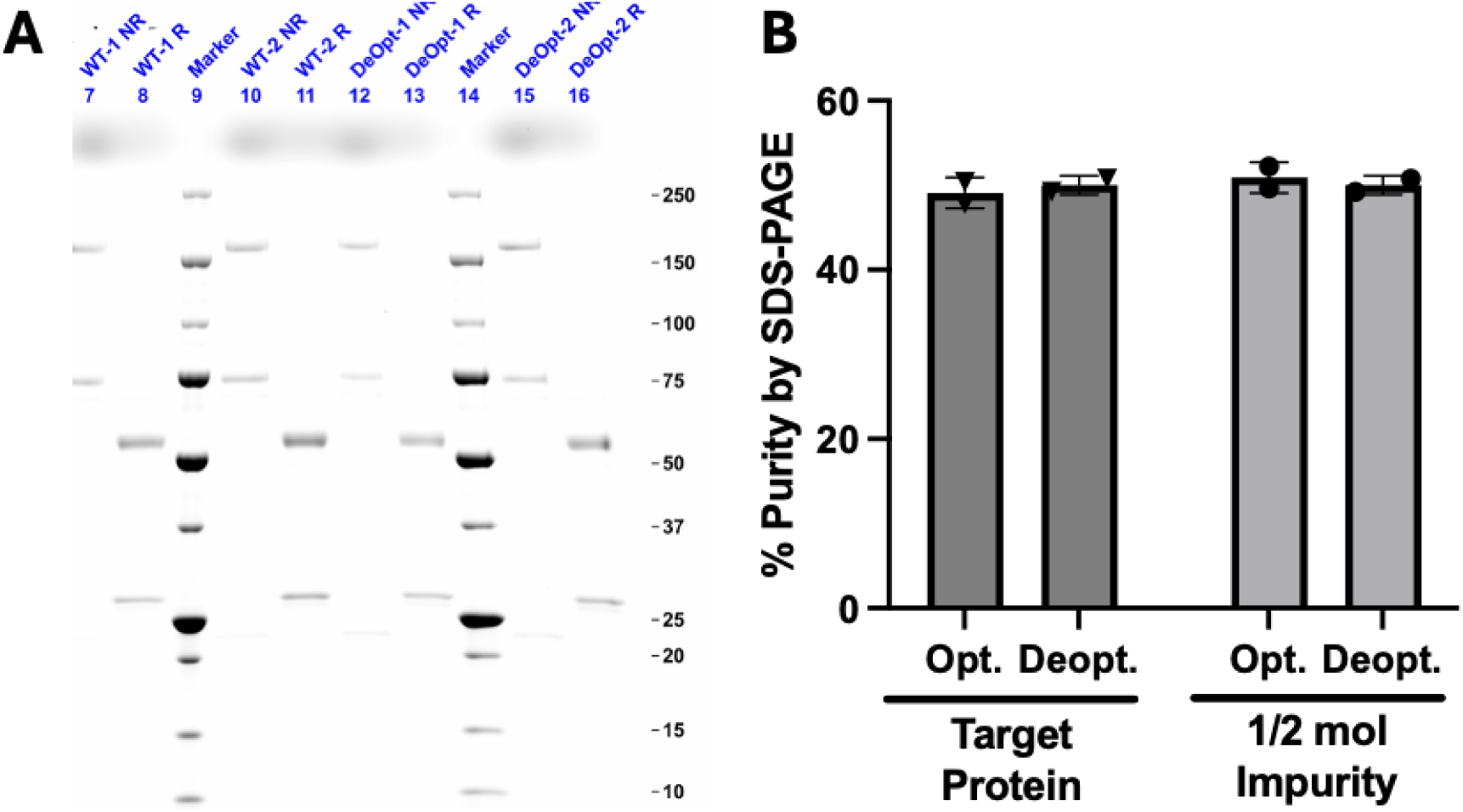
(A) Non-reduced and reduced SDS-PAGE of conditioned media (CM) from optimized (WT) and deoptimized expressions of Protein 2. (B) SDS-PAGE band quantitation of the target protein and half molecule bands in optimized and deoptimized expressions.

## References

1. Ellerman DA. The Evolving Applications of Bispecific Antibodies: Reaping the Harvest of Early Sowing and Planting New Seeds. BioDrugs. 2025;39(1):75–102.

2. Madsen AV, Pedersen LE, Kristensen P, Goletz S. Design and engineering of bispecific antibodies: insights and practical considerations. Front Bioeng Biotechnol. 2024;12:1352014.

3. Carter P. Bispecific human IgG by design. J Immunol Methods. 2001;248(1-2):7–15.

4. Krah S, Sellmann C, Rhiel L, Schroter C, Dickgiesser S, Beck J, et al. Engineering bispecific antibodies with defined chain pairing. N Biotechnol. 2017;39(Pt B):167–73.

5. Estes B, Sudom A, Gong D, Whittington DA, Li V, Mohr C, et al. Next generation Fc scaffold for multispecific antibodies. iScience. 2021;24(12):103447.

6. Merchant AM, Zhu Z, Yuan JQ, Goddard A, Adams CW, Presta LG, et al. An efficient route to human bispecific IgG. Nat Biotechnol. 1998;16(7):677–81.

7. Gong S, Wu C. Efficient production of bispecific antibodies—optimization of transfection strategy leads to high-level stable cell line generation of a Fabs-in-tandem immunoglobin. Antibody Therapeutics. 2023;6(3):170–9.

8. Li J, Menzel C, Meier D, Zhang C, Dubel S, Jostock T. A comparative study of different vector designs for the mammalian expression of recombinant IgG antibodies. J Immunol Methods. 2007;318(1-2):113–24.

9. Ayyar BV, Arora S, Ravi SS. Optimizing antibody expression: The nuts and bolts. Methods. 2017;116:51–62.

10. Rosenblum G, Chen C, Kaur J, Cui X, Zhang H, Asahara H, et al. Quantifying elongation rhythm during full-length protein synthesis. J Am Chem Soc. 2013;135(30):11322–9.

11. Yan X, Hoek TA, Vale RD, Tanenbaum ME. Dynamics of Translation of Single mRNA Molecules In Vivo. Cell. 2016;165(4):976–89.

12. Purvis IJ, Bettany AJ, Santiago TC, Coggins JR, Duncan K, Eason R, et al. The efficiency of folding of some proteins is increased by controlled rates of translation in vivo. A hypothesis. J Mol Biol. 1987;193(2):413–7.

13. Thanaraj TA, Argos P. Protein secondary structural types are differentially coded on messenger RNA. Protein Sci. 1996;5(10):1973–83.

14. Komar AA, Lesnik T, Reiss C. Synonymous codon substitutions affect ribosome traffic and protein folding during in vitro translation. FEBS Lett. 1999;462(3):387–91.

15. Goncalves-Carneiro D, Bieniasz PD. Mechanisms of Attenuation by Genetic Recoding of Viruses. mBio. 2021;12(1).

16. Coleman JR, Papamichail D, Skiena S, Futcher B, Wimmer E, Mueller S. Virus Attenuation by Genome-Scale Changes in Codon Pair Bias. Science. 2008;320(5884):1784–7.

17. Groenke N, Trimpert J, Merz S, Conradie AM, Wyler E, Zhang H, et al. Mechanism of Virus Attenuation by Codon Pair Deoptimization. Cell Rep. 2020;31(4):107586.

18. Sato S, Ward CL, Kopito RR. Cotranslational ubiquitination of cystic fibrosis transmembrane conductance regulator in vitro. J Biol Chem. 1998;273(13):7189–92.

19. Zhou M, Fisher EA, Ginsberg HN. Regulated Co-translational ubiquitination of apolipoprotein B100. A new paradigm for proteasomal degradation of a secretory protein. J Biol Chem. 1998;273(38):24649–53.

20. Quax TE, Claassens NJ, Soll D, van der Oost J. Codon Bias as a Means to Fine-Tune Gene Expression. Mol Cell. 2015;59(2):149–61.

21. Alexaki A, Hettiarachchi GK, Athey JC, Katneni UK, Simhadri V, Hamasaki-Katagiri N, et al. Effects of codon optimization on coagulation factor IX translation and structure: Implications for protein and gene therapies. Sci Rep. 2019;9(1):15449.

22. Komar AA. A pause for thought along the co-translational folding pathway. Trends Biochem Sci. 2009;34(1):16–24.

23. Kumada Y, Sakan Y, Kajihara H, Kihara M, Kikuchi Y, Yamaji H, et al. Efficient production of single-chain Fv antibody possessing rare codon linkers in fed-batch fermentation. J Biosci Bioeng. 2009;107(1):73–7.

24. Magistrelli G, Poitevin Y, Schlosser F, Pontini G, Malinge P, Josserand S, et al. Optimizing assembly and production of native bispecific antibodies by codon de-optimization. MAbs. 2017;9(2):231–9.

25. Steinmetz A, Vallée F, Beil C, Lange C, Baurin N, Beninga J, et al. CODV-Ig, a universal bispecific tetravalent and multifunctional immunoglobulin format for medical applications. MAbs. 2016;8(5):867–78.

26. Chen SW, Zhang W. Current trends and challenges in the downstream purification of bispecific antibodies. Antib Ther. 2021;4(2):73–88.

